# Directional invariance in the *Drosophila* giant fiber escape circuit

**DOI:** 10.1101/2022.07.15.500271

**Authors:** HyoJong Jang, David P Goodman, Catherine R von Reyn

## Abstract

An animal’s nervous system recognizes visual features regardless of where the visual feature is located. However, the underlying mechanisms that enable location invariant feature detection are not fully understood, particularly when visual feature information contributed by each eye needs to be resolved. Here we investigate directional invariance of looming responses in descending neurons (DN) of *Drosophila melanogaster*. We find multiple looming responsive DN integrate looming information across both eyes, even though their dendrites are unilateral. One DN in particular, the giant fibers (GF), generates invariant looming responses across approach directions. We confirm visual information propagates to the GF from the contralateral eye through an as of yet unidentified pathway and demonstrate the absence of this pathway alters GF responses to looming stimuli from the ipsilateral eye. Our data highlight a role for bilateral visual integration in generating consistent escape responses that are robust across a wide range of stimulus locations and parameters.

## Introduction

In responding to a visual stimulus, an animal’s nervous system needs to be able to deal with where the stimulus is located in its environment. In some cases, the stimulus location provides information to select a behavioral response (Bianco et al., 2011; Gancedo et al., 2020; Pouget et al., 2000). In other cases, the behavioral response needs to be invariant to stimulus location (Benucci, 2022; DiCarlo et al., 2012; Quiroga et al., 2005; Rolls, 2012). Although we know, to some extent, how directional information guides responses (Allen et al., 2021; Peek, 2018; Pouget et al., 2000), the underlying mechanisms that enable location invariance for sensory to motor transformations are not fully understood as the neural circuits that bridge sensory encoding and the selection of an appropriate behavior are largely incomplete (DiCarlo et al., 2012; Quiroga et al., 2005; Rolls, 2012).

This is well illustrated in visually evoked escape behaviors. Animals, when responding to an object approaching on a direct collision course, select amongst multiple potential behavioral responses. Animals may freeze, flee, or (for those that have the option) may takeoff into the air (Branco et al., 2020; Evans et al., 2019; Zacarias et al., 2018). Additional flexibility is witnessed within each response. For example, individual responses can consist of a flexible sequence of behaviors, like combining freezing with fleeing, or preparatory behaviors like postural adjustments that establish takeoff trajectories (Branco et al., 2020; Card et al., 2008; De Franceschi et al., 2016). Certain behaviors depend on the location of the stimulus. The fleeing direction of crabs and mice, the postural adjustments that precede the takeoff of the fruit fly *Drosophila melanogaster*, and the body bends of the C-start in telost fish are highly biased by the stimulus location (Bhattacharyya et al., 2017; Dunn et al., 2016; Fetcho et al., 1998; Hemmi et al., 2012; Korn et al., 2005). Alternatively, some behaviors appear to be invariant of stimulus location. For example, certain freezing behaviors and the leg extension and wing depression that comprise the takeoff jump of *Drosophila* occur regardless of stimulus location (Card et al., 2008; von Reyn et al., 2014; Zacarias et al., 2018).

Outside of the directional C-start responses of telost fish (Domenici et al., 2019), we know very little about the underlying circuit mechanisms for resolving the location of an object approaching on a direct collision course - in particular, how ipsilateral, contralateral or bilateral information are integrated to inform escape decisions. Looming stimuli (the 2D projection of an approaching object) have been used to identify potential collision detecting neurons across species (Dunn et al., 2016; Fotowat et al., 2011; Gabbiani et al., 1999; Liu et al., 2011; Oliva et al., 2014; Sun et al., 1998; von Reyn et al., 2014), and receptive field mapping has been performed on looming responsive retinal ganglion cells (RGC) and visual projections neurons (VPN) in the vertebrate retina and invertebrate optic lobe, respectively (Kerschensteiner, 2022; Klapoetke. et al., 2022; Morimoto et al., 2020). But limited investigations have been conducted at the next level of processing, where the information RGC and VPN provide from each eye are then combined in the brain. In vertebrates, this is predicted to first occur within the superior colliculus/optic tectum (Isa et al., 2021; Lee et al., 2020; Temizer et al., 2015; Zhao et al., 2014). In invertebrates such as the fruit fly, this is predicted to first occur in the optic glomeruli where a population of descending neurons (DN) receive information from VPN and then descend into the ventral nerve cord of the fly to synapse on motor and premotor circuits (Namiki. et al., 2018; Wu et al., 2016).

Here we investigate directional dependencies of looming responses in DN predicted to participate in looming evoked behaviors. We surprisingly find multiple looming responsive DN have significant responses to contralateral looming stimuli, even though their dendrites are ipsilateral. Focusing on one DN, called the giant fibers (GF), we find responses are invariant across object approach directions. Occluding one eye with paint, we demonstrate visual information propagates from the contralateral eye through an as of yet unidentified pathway, and that this information is key for the GF response to be maintained regardless of direction. We also find integration across both eyes is necessary for maintaining the GF response across loom speeds.

## Materials and Methods

### Fly Stocks

*Drosophila melanogaster* were raised on standard cornmeal fly food at 25°C and 50% humidity on a 12-h light-dark cycle. Female *Drosophila*, due to their larger size, were used for all electrophysiology, anatomy, and behavioral experiments 2–5 days after eclosion. All information pertaining to fly stocks is contained in Table 1.

**Table 1.** Drosophila stocks.

| Fly Stock | Source | IDENTIFIER |
| --- | --- | --- |
| GF-split-GAL4: (68A06_AD (attP40); 17A04_DBD (attP2)) | (von Reyn et al., 2014) | N/A |
| DNp02-split-GAL4: (VT063736_AD (attP40); VT017647_DBD (attP2)) | (Namiki. et al., 2018) | splitgal4.janelia.org: SS01554 |
| DNp03-split-GAL4: (R30C12_AD (attP40); R22D06_DBD | (Namiki. et al., 2018) | splitgal4.janelia.org: SS01066 |
| DNp06-split-Gal4: (*VT019018_AD (attp40); VT017411_DBD (attp2)*) | (Namiki. et al., 2018) | splitgal4.janelia.org: SS02256 |
| *w+ UAS-GFP* | (Pfeiffer et al., 2010) | N/A |

### Electrophysiology

In-vivo whole-cell, current-clamp electrophysiology was carried out on behaving, tethered flies as previously described (von Reyn et al., 2017). Flies were anesthetized at 4 °C and their head and thorax were tethered to polyether ether ketone (PEEK) plates with UV glue. The T1 legs were cut at the femur to avoid cleaning of the head and occlusion of the eyes. The proboscis was glued in its retracted position to decrease brain movement during the recording. The antennae were also immobilized with UV glue to limit stimulation of antennal afferents to the GF. For eye painting experiments, black paint (Golden High Flow Paint, carbon black acrylic) was then applied to one eye. To access GF for recordings, the cuticle and trachea on the posterior side of the head above the GF soma were removed and the brain was perfused with standard extracellular saline (NaCl 103 mM, KCl 3 mM, TES 5 mM, Trehalose-2H2O 8 mM, Glucose 10 mM, NaHCO3 26 mM, NaH2PO4 1 mM, CaCl2 - 2H2O 1.5 mM, MgCl2 - 6H2O 4 mM (Gouwens et al., 2009)) with the osmolarity adjusted to 270-275 mOsm and bubbled with 95% O_2_ / 5% CO_2_ to maintain a pH of 7.3. All experiments were performed at room temperature (20 °C - 22 °C). To access soma for recordings, a brief, localized application of collagenase (0.5% in extracellular saline) with a glass electrode was used to disrupt the brain sheath. GFP labeled soma were then targeted by patch clamp electrodes (3-6 MΩ) containing intracellular saline (K-Aspartate 140 mM, KCl 1 mM, HEPES 10 mM, EGTA 1 mM, Na3GTP 0.5 mM, MgATP 4 mM, Alexafluor-568 5 µM, 265 mOsm, pH 7.3). In-vivo whole-cell recordings were conducted in current-clamp mode using a MultiClamp 700B amplifier, and digitized (NI-DAQ, National Instruments) at 20 kHz. All data were acquired using the opensource software Wavesurfer (https://www.janelia.org/open-science/wavesurfer) running in Matlab. Traces were not corrected for a 13mV liquid junction potential (Gouwens et al., 2009). Recordings were considered acceptable when the initial seal resistance was >2 GΩ before rupture, the resting membrane potential was less than -50 mV, and the input resistance was > 50 MΩ.

### Visual Stimuli for Electrophysiology

Visual stimuli were back-projected onto a 4.5-inch diameter mylar cylindrical screen covering 180° in azimuth via two DLP projectors (Texas Instruments Lightcrafter 4500). To project onto a cylindrical surface, each projector was calibrated as described previously (Goodman et al., 2018) and an 18° overlap between the two projectors was blended for uniform illumination (Fig 1C). Following calibration and blending, each 912 × 1140 resolution projected image was displayed in 6bit grayscale at 240 Hz, above the flicker fusion frequency of *Drosophila* (100 Hz, (Niven et al., 2003)). All stimulus frames were generated in MATLAB and presented using psychtoolbox (http://psychtoolbox.org/, Brainard 1997).

**Figure 1.**
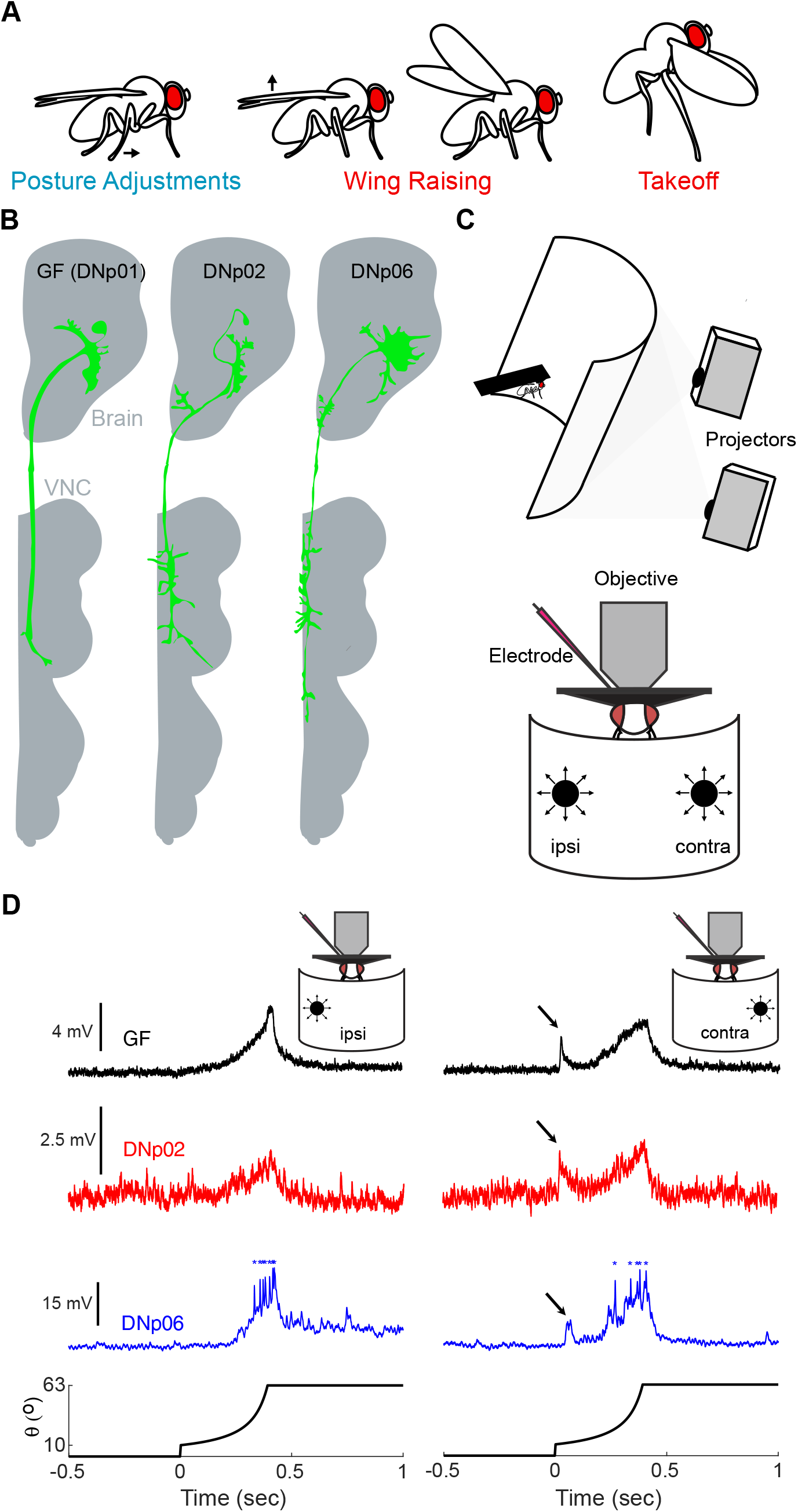
Multiple looming responsive DN integrate looming information both ipsilateral and contralateral to their dendrite locations. (**A**) Certain sub-behaviors depend on the location of a looming stimulus in the fly’s visual field, including the type of postural adjustment necessary to set the fly’s escape trajectory in a direction away from the stimulus (Card et al., 2008). Other sub-behaviors, including wing raising, wing depression and leg extension, occur across all stimulus directions (Card et al., 2008). (**B**) The anatomy of three putative looming responsive DN. Notice all dendrites in the brain are restricted to the hemisphere ipsilateral to the soma. (**C**) Head fixed fly recording preparation. Looming stimuli are displayed on a cylindrical screen while DN activity is recorded using whole-cell patch clamp electrophysiology. (**D**) Ipsilateral and contralateral looming responses of the three DN in (B). GF and DNp02 are average responses across flies (n=5 and 2, respectively). DNp06 is a single trial response, with asterisks indicating spikes. Bottom trace shows the angular size of the looming stimulus over time (*r*/*v* = 40 ms). Arrows indicate responses to the stimulus pop on.

Looming stimuli are the 2D projection of an object approaching at a constant velocity. The angular size (*θ*) of the stimulus subtended by the approaching object can be calculated in time (*t*) by the following equation (Gabbiani et al., 1999):

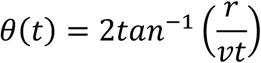

where *t* < 0 before collision and *t* = 0 at collision for an approaching object with a half size (*r*) and constant velocity (*ν*). As described previously (Gabbiani et al., 1999), *r*/*ν* (ms) is the half size to approach speed ratio. The smaller the *r*/*v*, the more abrupt the stimulus expansion.

The entire set of visual stimuli consisted of four looming stimuli (*r*/*v*=: 10, 20, 40, 80ms) starting at 10° and expanding to 63° and then held for 1 second, three constant angular velocity stimuli (100°/s, 2000°/s 6000°/s) starting at 10° and expanding to 63° and then held for 1 second, and small object motion stimuli (25°/s moving from left to right). All stimuli were displayed at three locations (45°, 0, -45° in azimuth, 0° in elevation). Each visual stimulus was presented once per trial, in a randomized order, at 30 second intervals. Two full trials were then averaged per individual fly.

### Direction selectivity

A direction selectivity index (DSI) was calculated for each fly across stimulus locations to see if the GF in individual flies was more selective to ipsilateral stimuli over contralateral stimuli or to ipsilateral stimuli over center stimuli (Elstrott et al., 2008; Hei et al., 2014):

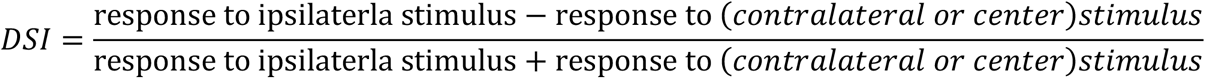

To evaluate whether DSI scores indicated significant direction selectivity, a permutation test (Baden et al., 2016) was performed by generating a null hypothesis distribution (10000 shuffled trials for responses between the two direction) representing no directional bias. The p-value was then computed by calculating the percentage of the null distribution above the mean of the group DSI.

### Data Analysis and Statistics

All analyses were performed using in house MATLAB scripts. Electrophysiology recordings for each stimulus presentation were baseline subtracted by taking the average response one second preceding the stimulus onset. The GF expansion peak magnitude and latency (following stimulus onset) during loom expansion were measured after filtering each recording (Savitzky-Golay, 4th order polynomial, frame size is 1/10th the length of the stimulus). The pop on peak magnitude and latency were measured within a 50 ms time window after the onset of the stimulus. To select the appropriate parametric or non-parametric test, the Shapiro-Wilk test was used to evaluate whether data were normally distributed. All statistical tests applied to data are as stated in the figure captions.

## Results and Discussion

*Drosophila* responses to looming stimuli are both complex and flexible. They can consist of a sequence of motor programs (Fig. 1A), such as freezing, postural adjustments, and wing raising that precede the wing depression and leg extension of a takeoff. Additionally, each motor program may or may not be present in any escape, like freezing without takeoffs, or takeoffs without wing raising (von Reyn et al., 2014; Zacarias et al., 2018). The direction of approach of a looming stimulus impacts the selection of certain motor programs, like which postural adjustment needs to be made to establish takeoff trajectory (Card et al., 2008; Peek, 2018) (Fig 1A). Other motor programs, like the leg extension and wing depression of a takeoff, can be observed regardless of stimulus direction (Card et al., 2008; von Reyn et al., 2014).

The selection of escape motor programs is thought to happen at the level of descending neurons (DN) that integrate sensory information in the brain and relay this information down to premotor and motor centers in the *Drosophila* ventral nerve cord (spinal cord homologue). To date, only one DN the giant fibers (GF) have a well-established role in escapes, driving a leg extension and wing depression program for takeoffs (Ache et al., 2019; von Reyn et al., 2014; von Reyn et al., 2017). However, anatomical, functional and behavioral evidence have recently emerged that highlight additional DN may drive other motor programs witnessed in the behavior, or drive escapes in the absence of GF activation (Fotowat et al., 2009; Namiki. et al., 2018; Peek, 2018; von Reyn et al., 2014; Zacarias et al., 2018).

We first investigated the directional selectivity of looming responses of the GF and two other DN (DNp02 and DNp06) identified as looming-evoked escape behavior candidates (Fig 1B) (Namiki. et al., 2018; Peek, 2018). All three DN have dendrites located in one hemisphere of the central brain, ipsilateral to their soma (here on referred to as the ipsilateral side) and project axons down to the same side of the VNC. Based on this anatomy, we anticipated we would witness large DN responses to looming stimuli on the ipsilateral side, but significantly reduced responses on the contralateral side. We modified our previously published visual setup (Goodman et al., 2018) to expand our coverage of the fly’s frontal visual field to 180° (Figure 1C). We then recorded from head fixed animals, using whole-cell electrophysiology, the DN responses to looming stimuli with ipsilateral or contralateral approach directions (± 45° azimuth), with a dark disk size starting at 10° and expanding to 63° to limit the contribution from the eye opposite to the stimulus location (Fig 1C). Interestingly, we found all three DN displayed significant responses to both ipsilateral and contralateral visual stimuli, despite having unilateral dendrites (Fig 1D). These data suggest all three DN receive information from the contralateral eye through yet to be identified commissural pathways. We also found, as reported previously (Peek, 2018), that the DNs have different types of responses to these stimuli. For example, DNp02’s response remained subthreshold while DNp06’s response showed the largest depolarization with spikes appearing towards the end of the stimulus expansion.

We next expanded our stimulus subset to perform a deeper investigation into directional dependence by adding looming presentations from the front of the fly that would expand across both eyes, and concurrent ipsilateral and contralateral looming stimuli that would excite both eyes (Fig 2A). We also presented our looms at a range of radius/approach speed (*r/v*) values (Gabbiani et al., 1999), with small values indicating abrupt expansions and large values indicating more gradual expansions. We focused on responses in the GF.

**Figure 2.**
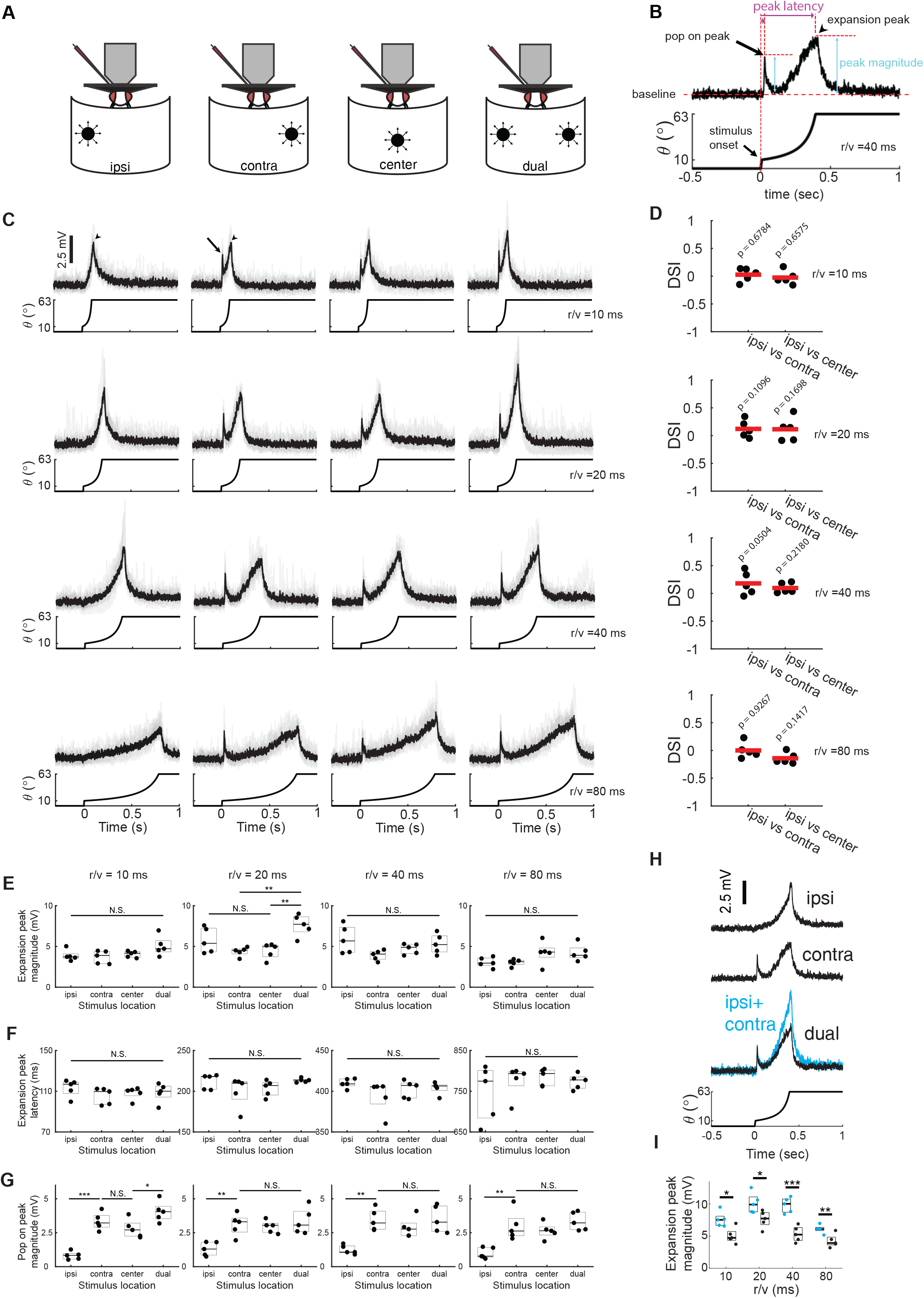
GF directional invariance to looming stimuli. (**A**) Four looming stimuli locations used for these investigations. (**B**) Measured components of a GF looming response (**C**) GF average (black) and per fly (gray) responses to looming stimuli across *r*/*v* and across location. (**D**) Direction selective index (DSI) comparing responses to ipsilateral vs contralateral stimuli and ipsilateral vs center stimuli. Statistical significance of DSI score was performed using a permutation test and p-values are displayed in each comparison. (**E**-**G**) Quantification of the data in *(C)* following the measurements in *(B)* (ANOVA, Tukey’s HSD post hoc, *=p<0.05, **=p<0.01, ***=p<0.001, n = 5 flies) (**H**) Example sum of a fly’s response to ipsilateral and contralateral stimuli as compared to the actual response to a dual stimulus (**I**) Quantification of data represented in *(H)* across all *r*/*v*. (ANOVA, Tukey’s HSD post hoc, *=p<.05, **p<.01, ***<.001, n=5 flies,).

To compare GF responses across this range of stimulus parameters, we measured components of the responses as indicated in (Fig 2B). We found the peak magnitude of the GF response during looming stimulus presentations was not significantly different across all stimulus locations (Fig 2C and 2E), ipsi, contra, and center, suggesting the GF is invariant to stimulus direction within the area of the visual field we evaluated. To investigate this further, we next calculated a direction selectivity index (DSI) for each fly across stimulus locations to see if the GF in individual flies was more selective to ipsilateral stimuli over contralateral stimuli or to ipsilateral stimuli over center stimuli (Fig 2D). While we found the DSI to be highly variable across flies, no DSI exceeded the 0.5 threshold to be considered directionally selective (Elstrott et al., 2008; Hei et al., 2014). Additionally, we found measured DSI were not significantly different from a randomized distribution of responses(Fig 2D, p-values). These data suggest the GF has minimal to no directional preference across the stimulus locations we evaluated.

Furthermore, we found minimal to no increase in the GF response when stimulating both eyes with the dual looming stimulus. We had predicted, based on witnessed linear and supralinear integration of ipsilateral visual inputs (Ache et al., 2019; von Reyn et al., 2017), that the dual stimulus response would at least be a sum of the ipsilateral and contralateral stimulus responses (Fig 2H). However, we found the GF response magnitude to the dual stimuli to be significantly less than the sum of the individual responses across all *r/v* (Fig 2I), and for the majority of *r/v* not significantly different from a single looming stimulus presentation (Fig 2E), suggesting the GF may quickly reach an upper bound on its response through intrinsic or circuit level mechanisms.

We next investigated the timing of the peak GF response across the different stimulus locations, as response timing between different looming DNs is hypothesized to guide escape behavior selection (Card et al., 2008; von Reyn et al., 2014). We hypothesized that responses emerging from the contralateral side may be delayed with respect to responses emerging from the ipsilateral side through known direct connections with VPN outputs from the optic lobe (Ache et al., 2019; von Reyn et al., 2017). However we found no significant different in the peak latency across directions (Fig 2F). As GF mediated takeoff responses follow a size threshold model (von Reyn et al., 2014), these data suggest the GF is able to consistently time its response based on an expanding stimulus’s size, regardless of the stimulus approach direction.

While the peak response magnitude and timing remained consistent across stimulus locations, we did witness a difference in the presence of a transient response that occurs when the looming stimulus first appears on the screen as a 10° disk. This “pop on” transient was present for contralateral looms but not for ipsilateral looms (Fig 2C), as evident by a significantly lower magnitude across flies (Fig 2G). Interestingly, this difference in pop on transient was also observed across other DNs (Fig 1D arrows). We hypothesized that the absence of the pop on transient in this investigation may be due to a difference in visual displays (Baden et al., 2016), as this transient occurs in GF during stimulus presentations to a single eye (Ache et al., 2019; von Reyn et al., 2017) while our display illuminates both eyes. Prior work has demonstrated that LC4 contributes predominantly to the pop on response in the GF (von Reyn et al., 2017). Our data therefore suggest brightness differences across eyes may modulate the LC4 contribution to DN responses.

Although our recordings from the GF strongly suggest the GF receives information from the contralateral eye, we wanted to validate these findings with eye occlusion experiments. We painted the contralateral eye of the fly with an opaque carbon black acrylic paint and displayed the same set of visual stimuli while recording from the GF. As expected, we found occlusion of the contralateral eye eliminated the GF response for all but the final expansion of the contralateral looming stimulus (Fig 3C, D, F). We assume the small response at the end of expansion marks when the stimulus crosses into the receptive field of the ipsilateral eye (Fig 3C, asterisk), within the fly’s binocular region (Buchner, 1971). These data confirm the GF receives substantial visual inputs from the contralateral eye. As the VPN LC4 and LPLC2 that provide ipsilateral visual input to the GF have small, ipsilaterally restricted receptive fields (Klapoetke. et al., 2022), these data suggest the GF receives contralateral information through an as of yet unidentified commissural pathway.

**Figure 3.**
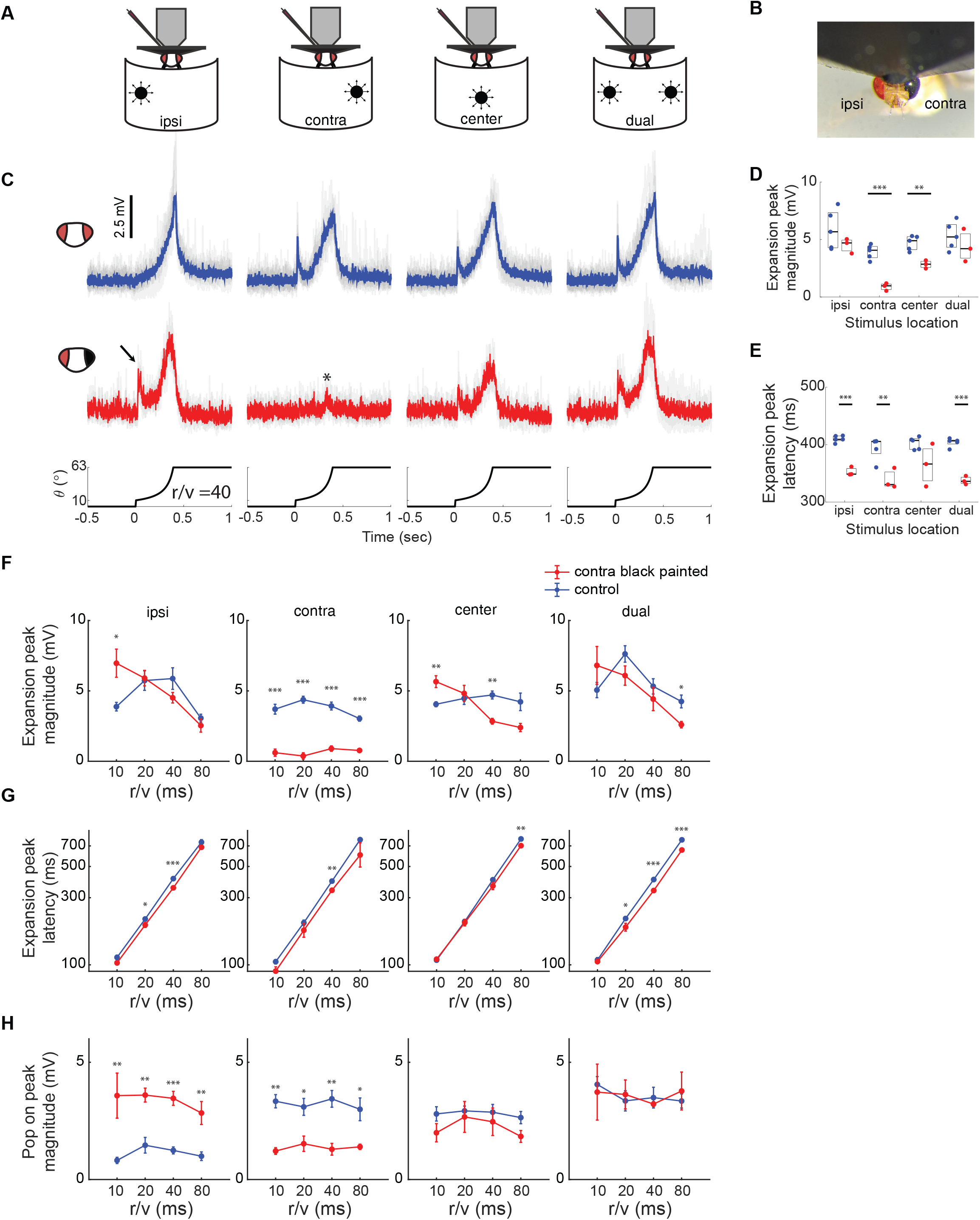
GF response tuning to ipsilateral looming stimuli is modulated by information from the contralateral eye. (**A**) Four looming stimuli locations used for these investigations. (**B**) Example contralateral eye painted, tethered fly. (**C**) Example GF average and per fly (gray) responses to looming stimuli (*r*/*v=*40ms) and across locations with (red) and without (blue) contralateral eye painting. The arrow indicates the pop on peak rescued and the asterisk indicates the response due to stimulation of the ipsilateral eye. (**D** - **E**) Quantification of the data in *(C)* (ANOVA, Tukey’s HSD post hoc, *=p<.05, **=p<.01, ***=p<.001, control n=5 flies, black painted n=3 flies). (**F-H**) expanded quantification of eye painting experiments across *r*/*v*, following the measurements in Figure 2B, values are displayed as mean ± SEM for control (blue) and eye painted (red) flies, (ANOVA, Tukey’s HSD post hoc, *=p<.05, **=p<.01, ***=p<.001, control n=5 flies, black painted n=3 flies).

We next investigated how the removal of contralateral information affected the GF responses to center and ipsilateral looming stimuli. We predicted contralateral eye painting would result in a partial reduction in GF peak response magnitude to stimuli expanding from the center, across both eyes. We anticipated eye painting to have minimal effects on the response to ipsilateral looms, at most causing a small reduction in the response near the end of the expansion, when the stimulus crosses into the receptive field of the contralateral eye. Although we did witness our anticipated findings of a reduction in the response to center stimuli and no change in the response to ipsilateral looms, it only occurred for larger *r*/*v* (Fig 3D,F). Unexpectedly, for ipsilateral and center stimuli, we witnessed enhanced responses to looms with an *r*/*v* of 10ms that shifted the GF’s tuning preference towards rapidly expanding stimuli. Our data suggest visual information from both eyes contributes to GF tuning. As LC4’s angular velocity contribution to the GF response magnitude increases as *r*/*v* decreases (while LPLC2’s angular size contribution remains constant), these data again suggest LC4 input to the GF circuit is particularly susceptible to brightness differences across both eyes.

We then investigated how contralateral eye painting affects the timing of the GF peak response (Fig 3G). We hypothesized that the loss of excitation from the other eye would decrease the latency of the peak GF ipsilateral, contralateral, and dual loom responses as they would no longer benefit from the stimulus expanding into the binocular region of the fly’s visual field at the end of the loom. In agreement with our hypothesis, we witnessed significant shifts to shorter latencies across these loom locations.

Finally, we investigated how contralateral eye painting changes the magnitude of the GF pop on peak response (Fig 3H). As the starting size of the stimulus does not change across *r*/*v*, we did not anticipate significant differences in magnitude except a complete loss of the response in the painted contralateral eye. While the contra, center and dual stimuli results followed our predictions, we unexpectedly regained the pop on transient in the ipsilateral eye. While unexpected, these results support our hypothesized reason the pop on peak is lost in our control flies: LC4 contributions are modulated by brightness differences across the eye.

In summary, we found a subset of DNs (GF-DNp01, DNp02, DNp06) respond to looming information from the contralateral eye even though their dendrites are ipsilateral (Namiki. et al., 2018). These data suggest yet to be identified commissural pathways provide contralateral looming information. We also observed each DN has a distinct looming response (with differences in magnitude, timing, and spiking). Since *Drosophila* escape responses are predicted to require multiple DN to enable behavioral complexity and flexibility (Namiki. et al., 2018; Peek, 2018; von Reyn et al., 2014; Zacarias et al., 2018), we assume the difference in looming response could be mapped onto different behaviors or sub-behaviors that comprise escape responses (Card et al., 2008).

We found one DN, the GF, generates responses that are invariant to looming stimulus direction. This invariance is aligned with the GF’s role in escape: the GF drive a synchronized wing depression and synchronized leg extension that are present in takeoff escapes regardless of the stimulus position (Allen et al., 2006; Card et al., 2008; von Reyn et al., 2014). The GF circuit’s ipsilateral connectivity is relatively well studied (Ache et al., 2019; von Reyn et al., 2017) while its contralateral connectivity is not well established. GF axons form electrical synapses in the VNC (Allen et al., 2006), however the similar latency and magnitude of contralateral and ipsilateral responses suggest contralateral visual information is not arriving from this connection that would be attenuated and delayed. The GF is also electrically coupled to giant commissural interneurons (also known as AMMC-A1) that receive direct information from the Johnson’s organ of the fly’s auditory system on the contralateral hemisphere (Pézier et al., 2016), but contralateral visual pathways have yet to be investigated.

While our work here focuses on ipsilateral and contralateral invariance, recent investigations have identified looming sensitive DNs (including DNp02) that may have anterior/posterior direction selectivity based on their VPN inputs and optogenetically elicited behaviors that establish forward or backwards escape trajectories (Peek, 2018). However, their directional tuning has yet to be investigated. Optogenetic activation of the GF showed no shift in body position or imposed directionality in escape trajectories (Peek, 2018), suggesting invariance may also exist on this axis.

Finally, we found the illumination level of the contralateral eye modifies the processing of dynamic stimuli in the ipsilateral eye. Occluding the contralateral eye enhanced ipsilateral tuning to abrupt stimuli – the appearance of a 10° disk or a rapidly expanding disk (Fig 3C, arrow). This may be similar to the luminance gain modulation or compensatory plasticity occurring in vertebrates following an occlusion of the contralateral eye (Ding et al., 2006; Dougherty et al., 2021; Lunghi et al., 2011; Moradi et al., 2009; Sato et al., 2008). Our work suggests this modulation has differential effects on visual features – a large effect on angular velocity (LC4) but a minimal to small effect on angular size (LPLC2) encoding. Interestingly, vertebrate occlusion studies have also witnessed differential effects across visual feature pathways (Lunghi et al., 2013). The underlying mechanisms however, are not well established. We believe future work in the *Drosophila* GF circuit will help shed light on these mechanisms.

## Acknowledgements

No

## Author contributions

HyoJong Jang (experimental design, stimulus design, electrophysiology, data analysis, wrote paper), David Goodman (stimulus design), Catherine R. von Reyn (conceptualization, experimental design, wrote paper)

